# Dynamical interactions reconfigure the gradient of cortical timescales

**DOI:** 10.1101/2020.10.23.350322

**Authors:** P Sorrentino, G Rabuffo, F Baselice, E Troisi Lopez, M Liparoti, M Quarantelli, G Sorrentino, C Bernard, V Jirsa

## Abstract

A hierarchy of local timescales with a back (sensory)-to-front (prefrontal) gradient reflects brain region specialization. However, cognitive processes emerge from the coordinated activity across regions, and the corresponding timescales should refer to the interactions rather than to regional activity. Using edgewise connectivity on magnetoencephalography signals, we demonstrate a reverse front-to-back gradient when non-local interactions are prominent. Thus, the timescales are dynamic and reconfigure between back-to-front and front-to-back patterns.

## Introduction

Conceptualizing the brain as a network has led to the identification of invariant features of large-scale activity (Sporns, 2013). In fact, brain activity displays structured topographies across different imaging modalities (e.g. resting state networks, gradient of timescales). In particular, brain regions (nodes) that are hierarchically lower in information processing operate at higher speed as compared to associative areas, which integrate information (Murray et al., 2014a, 2017;Gao et al., 2020; Müller et al., 2020; Shafiei et al., 2020). Such back-to-front gradient of timescales considers local information processing only. However, acquisition, integration and interpretation of inputs are distributed and dynamical processes, relying on the interactions (functional edges) occurring between regions. Hence, we hypothesize that the corresponding time-scales are inherent to the coordinated activity between regions, and not to local processing alone. So far, the hierarchical organization at the edge level is not known.

## Results

Using source-reconstructed magnetoencephalographic (MEG) data from a cohort of 58 healthy young subjects, we analyzed the time-resolved correlations between all pairs of brain regions, as a proxy of dynamical interactions. Namely, for each edge in the brain network we defined a time series as the co-fluctuations of the MEG signals at the extremal nodes (Fig.1.a) (See Materials and Methods). We used the Auto Mutual Information (AMI) to measure the amount of statistical dependence between any co-activation time series and its time-delayed copy (Mackay, 1995).

**Fig. 1.**
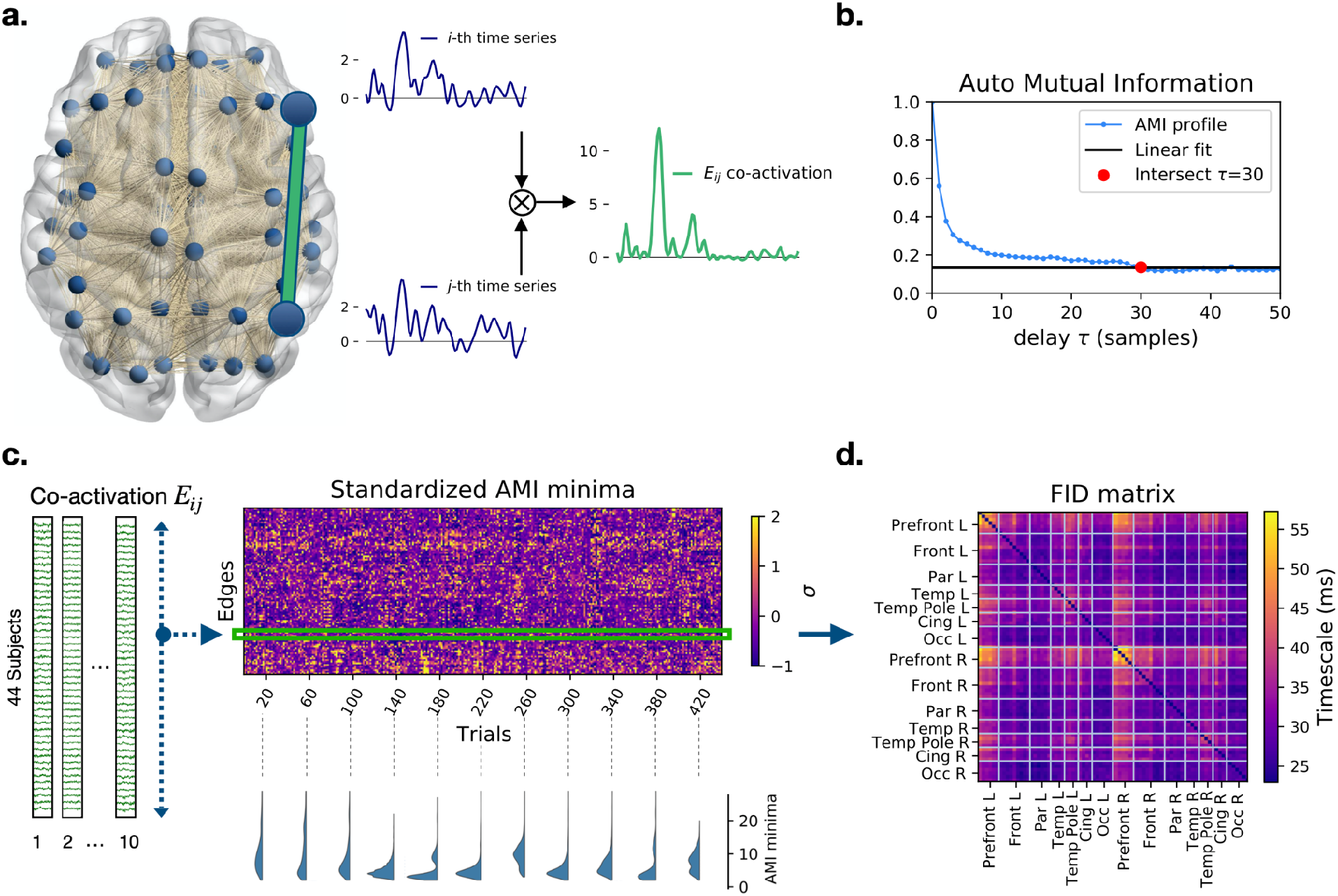
Auto mutual information analysis of the edge time-series in MEG data. a. The preprocessed magnetoencephalography (MEG) dataset consists of N = 78 source-reconstructed signals, one for each brain region. We define the edge time series E_ij_(t) (green) as the elementwise product of nodal z-scored signals at i and j (blue). b. Each point in the plot represents the mutual information between the co-activation time series and its τ-delayed copy. The red dot represents the delay at which the AMI reaches a minimal level (defined as a linear fit of the AMI tail). Notice that each time step τcorresponds to 3.9 milliseconds, given the sample frequency of 256 Hz. c. For each of the 44 subjects, the time series are split into 10, 10-second-long segments (epochs). For each trial, the τcorresponding to the AMI minima is computed. In the carpet plot, the minima are standardized across edges for each trial separately. The example minima distributions (bottom) in randomly selected epochs show that multimodality can emerge naturally d. Averaging over the standardized AMI minima, we obtain the N × N Functional

Repeating this operation for several delays, a profile of information decay was drawn for each edge of the brain network (Fig.1.b). For short time delays, the high value of the AMI indicates little information loss. The AMI drops (loss of information) as the time delay increases. A fast/ slow characteristic decay time indicates fast/slow information loss.

AMI edge decay times are organized according to a characteristic spatio-temporal pattern (Fig.1.c, top right). The trial-specific distributions of the decay times can be both multimodal and unimodal, showing that the brain can dynamically rearrange into sub-networks operating at different timescales (examples are shown in Fig.1.c, bottom right). Averaging across trials, we define the Functional Information Decay (FID) matrix (Fig.1.d), which reveals a temporal hierarchy of the edges. Considering the edges with the lowest and highest retention of information in the FID matrix, we identify two sub-networks: the short- and long-term storage network (SSN, Fig.2.a, left, LSN Fig.2.a, right, respectively). The SSN spans regions related to stimulus processing, while the LSN mainly involves regions related to higher cognitive functions (Table 1). We conclude that the hierarchy of timescales, consistent with the previously described nodal hierarchy, is manifest at the edge level (on average). Average information storage at the edge level ranges from 16 to 55 msec. However, at the single trial level, we find a larger range, with decays varying between 3.9 to 277 msec.

**Fig. 2.**
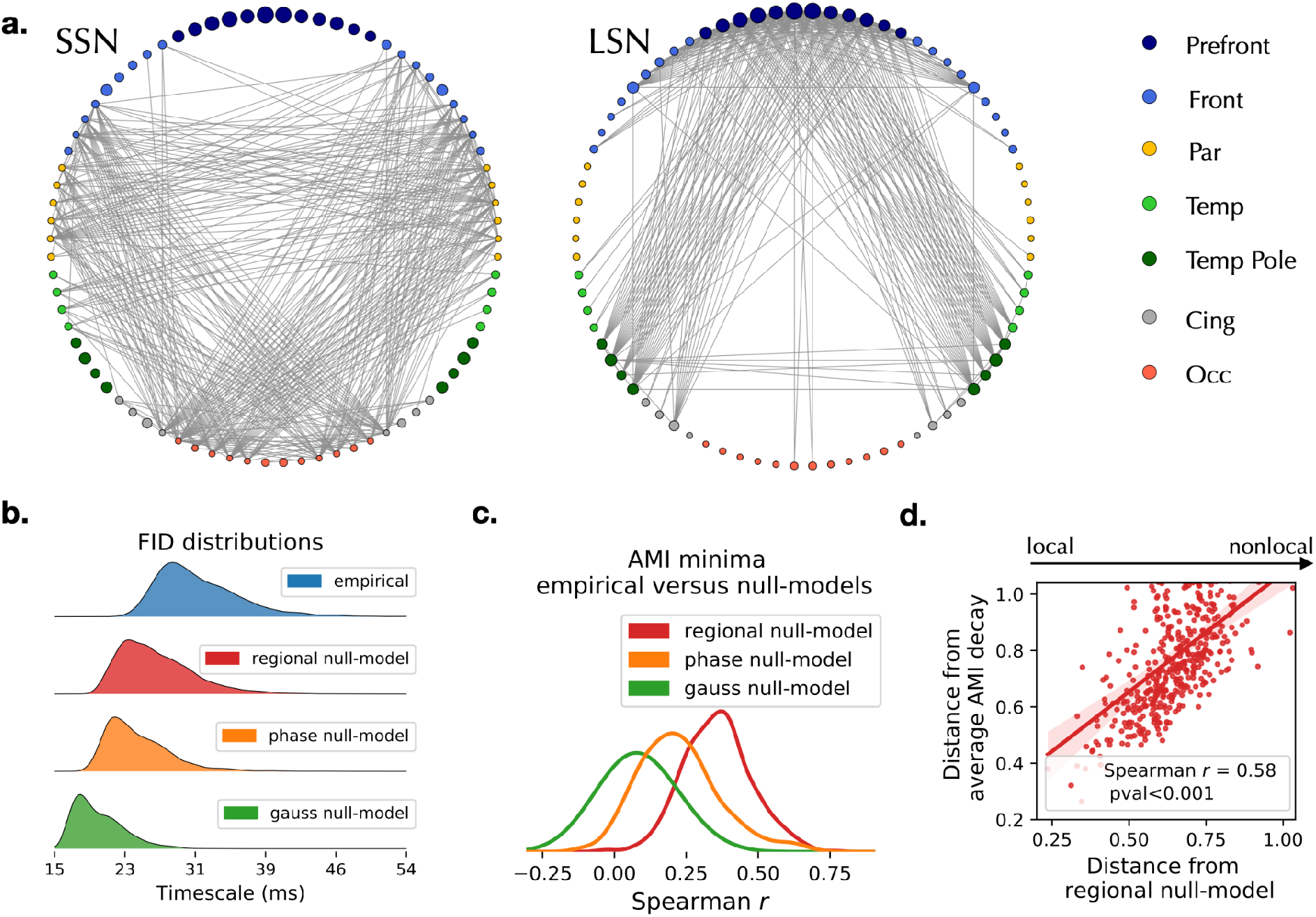
Topography of delays and surrogate analysis. a. The edges with the fastest (left) and slowest (right) decay times split into a short-storage and a long-storage network (SSN and LSN) at the trial-averaged level. b. Distributions of the average edge AMI decays in the original (blue), regional (red), phase (orange) and Gaussian (green) null-models show that the long decay times of information depend on the dynamics of the edges and are not explained by nodal spectral features or by static correlations alone. c. Distribution of the correlation between the decay times of each trial with the corresponding null-models. d. The x-axis measures the distance (1 - Spearman’s correlation) of each trial from the null-model, which is used to represent the amount of non-local interactions in each trial. The y-axis represents the distance from the FID matrix (averaged across empirical trials). Each dot represents a trial.

**Table 1.** List of regions of AAL Atlas. Each region number is colored according to the lobe. Color codes are reported in Fig.2.a.

| IDs | ROIs | IDs | ROIs |
| --- | --- | --- | --- |
| 1 | Left Rectus | 40 | Right Rectus |
| 2 | Left Olfactory | 41 | Right Olfactory |
| 3 | Left Frontal Sup Orb | 42 | Right Frontal Sup Orb |
| 4 | Left Frontal Med Orb | 43 | Right Frontal Med Orb |
| 5 | Left Frontal Mid Orb | 44 | Right Frontal Mid Orb |
| 6 | Left Frontal Inf Orb | 45 | Right Frontal Inf Orb |
| 7 | Left Frontal Sup | 46 | Right Frontal Sup |
| 8 | Left Frontal Mid | 47 | Right Frontal Mid |
| 9 | Left Frontal Inf Oper | 48 | Right Frontal Inf Oper |
| 10 | Left Frontal Inf Tri | 49 | Right Frontal Inf Tri |
| 11 | Left Frontal Sup Medial | 50 | Right Frontal Sup Medial |
| 12 | Left Supplementary Motor Area | 51 | Right Supplementary Motor Area |
| 13 | Left Paracentral Lobule | 52 | Right Paracentral Lobule |
| 14 | Left Precentral | 53 | Right Precentral |
| 15 | Left Rolandic Operculum | 54 | Right Rolandic Operculum |
| 16 | Left Postcentral | 55 | Right Postcentral |
| 17 | Left Parietal Sup | 56 | Right Parietal Sup |
| 18 | Left Parietal Inf | 57 | Right Parietal Inf |
| 19 | Left SupraMarginal | 58 | Right SupraMarginal |
| 20 | Left Angular | 59 | Right Angular |
| 21 | Left Precuneus | 60 | Right Precuneus |
| 22 | Left Heschl | 61 | Right Heschl |
| 23 | Left Temporal Sup | 62 | Right Temporal Sup |
| 24 | Left Temporal Mid | 63 | Right Temporal Mid |
| 25 | Left Temporal Inf | 64 | Right Temporal Inf |
| 26 | Left Temporal Pole Sup | 65 | Right Temporal Pole Sup |
| 27 | Left Temporal Pole Mid | 66 | Right Temporal Pole Mid |
| 28 | Left ParaHippocampal | 67 | Right ParaHippocampal |
| 29 | Left Fusiform | 68 | Right Fusiform |
| 30 | Left Cingulum Ant | 69 | Right Cingulum Ant |
| 31 | Left Cingulum Mid | 70 | Right Cingulum Mid |
| 32 | Left Cingulum Post | 71 | Right Cingulum Post |
| 33 | Left Insula | 72 | Right Insula |
| 34 | Left Occipital Sup | 73 | Right Occipital Sup |
| 35 | Left Occipital Mid | 74 | Right Occipital Mid |
| 36 | Left Occipital Inf | 75 | Right Occipital Inf |
| 37 | Left Calcarine | 76 | Right Calcarine |
| 38 | Left Cuneus | 77 | Right Cuneus |
| 39 | Left Lingual | 78 | Right Lingual |

Information Decay (FID) matrix, where rows and columns are regions, and the matrix elements are the average minima across trials. For each couple of brain regions this matrix describes the typical time that the information is preserved in the corresponding co-activation signal.

The average edge decay times vary smoothly across the brain, spanning between the SSN and the LSN (Fig.2.b, blue). Edge decay times may capture non-local interactions between regions or merely reflect local nodal processes. To distinguish between the two possibilities, we generate three null-models to disrupt or disregard dynamic interactions between regions: 1) gaussian (G-) surrogates, where the observed nodal autocorrelations (i.e., the power spectrum) are imposed on otherwise independent processes, 2) phase-randomized (P-) surrogates preserving both the regional autocorrelations and the pairwise static correlations (i.e., preserving static functional connectivity while disrupting the dynamics), and 3) regional null-models, where the edge decay times are defined as the geometric mean of the AMI decay times estimated from nodal timeseries 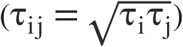. None of the three null-models reproduces the empirical decay-distribution (Fig.2.b). We conclude that edge time delays characterize dynamical interactions between regions.

To test whether the topographical organization of the edge decay times emerges from local or distributed processes, we correlate the empiric AMI decays with the decays retrieved by the null-models. At the trial-averaged level, the FID matrices derived from the null-models correlate with the empirical ones (Spearman’s r_s_ = 0.89, r_s_ = 0.94, r_*s*_ = 0.95, for the G-, P- and regional nullmodels, respectively, p < 0,001 for all cases). However, a greater variability of the correlations exists at the single-trial level (Fig.2.c), with trials that do not show significant correlations to the corresponding null-models. The regional null-model is the closest one to the empirical data, thus we selected it for further analyses. Since the regional null-model only retained local properties, we classified each empirical trial according to its distance from the corresponding null-model (defined as: 1 - the Spearman’s correlation coefficient in Fig.2d), Hence, a trial which is distant from its null-model is one that possess prominent non-local features. Notably, the more an empirical trial has non-local interactions, the more it is distant from the empirical (average) FID matrix (*r_s_* =0.58; Fig.2.d). To summarize, we have discovered a set of trials that possess significant non-local (edge) properties, which makes the topography of timescales deviate from the average configuration.

Strikingly, the corresponding trial in the null model recovers the empirical average topography (Fig.3a; delays are standardized along the edges within each trial). In fact, the correlation between the empirical trial and the empirical FID matrix is lower than the correlation between the corresponding surrogate trial and the empirical FID matrix (Fig.3.b, top left; Note that most correlations fall above the diagonal - represented by the orange line. Local to non-local trials are represented in colors from dark to light gray). Subtracting the standardized empirical delays from the null-model ones (Fig.3b, bottom), we show that the magnitude of the deviation from the null model is higher for non-local trials (Fig. 3b, top right). Averaging the deviations across trials, we show high correlation between the average FID matrix and the average deviation matrix (Fig. 3.c, bottom; r_s_ = 0.79). That is, the edges manifesting faster dynamics than expected from the regional null-model (negative average deviations) are roughly corresponding to the LSN, while slower-than-expected edges (positive average deviations) generally belong to the SSN (Fig. 3.c, top; edges are sorted according to trial-average delay. Color map as in Fig.1.d). We note that SSN edges are characterized by higher deviation variability as compared to the LSN ones.

**Fig. 3.**
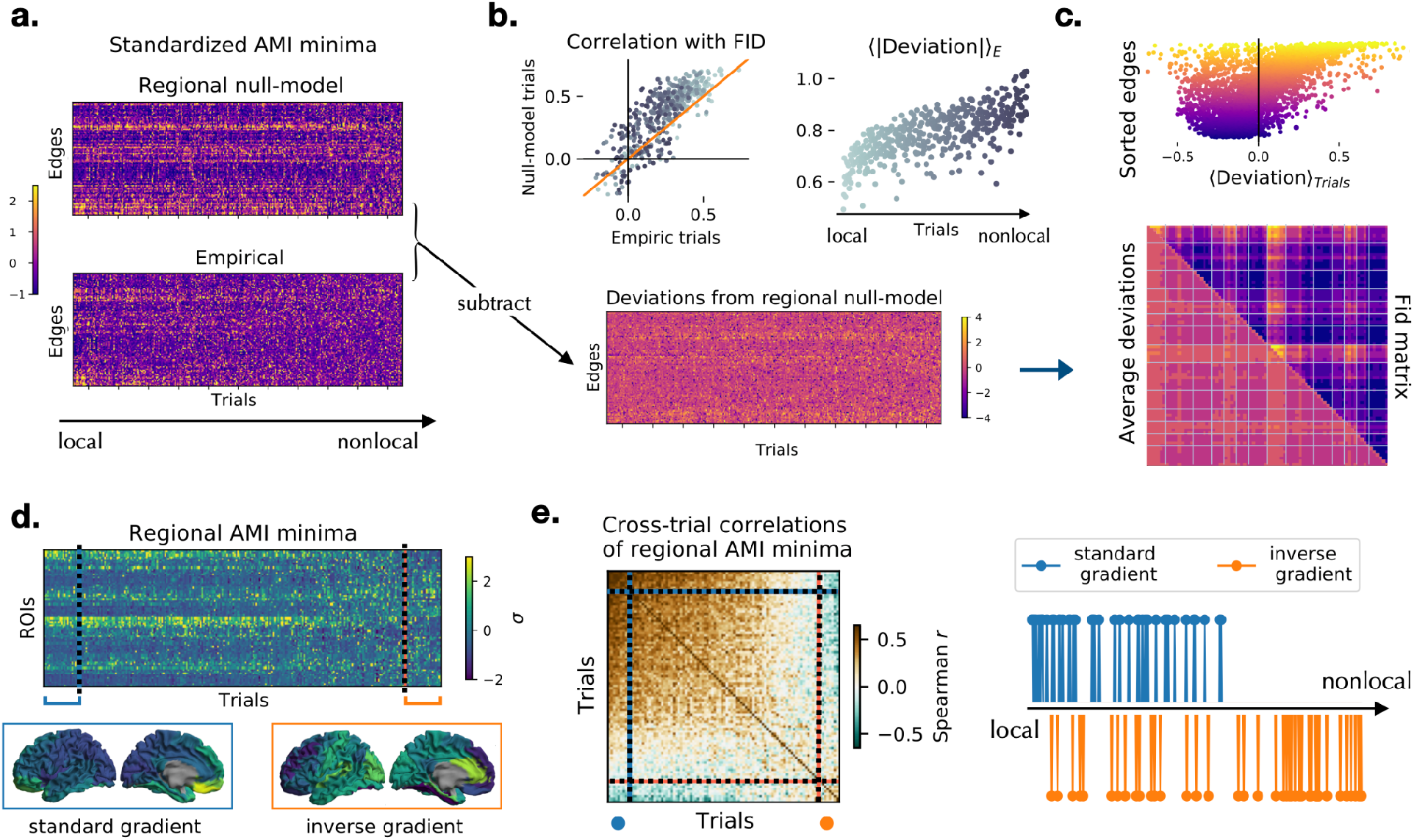
Trial-specific topography of edge and nodal timescales. a. Standardized edge decay times (AMI minima) of the regional null-model (top) and the empirical trials (bottom). The trials (matrix columns) are sorted according to a local–to–non-local axis, with the highest (lowest) correlation between the empirical and null-model trials in the first (last) columns. b, bottom. The empirical minima (panel a., bottom) are subtracted from the null-model minima (panel a., top) to define the trial-specific deviations from the null-model. b, top left. Correlation among the FID matrix (trial-average of the empirical AMI minima) and the trial-specific AMI minima in the empirical (x-axis) and null-model (y-axis). Most trials are above the diagonal (orange), showing that the null-model retrieves the empirical average topography of timescales better than the empirical single trials. Dots represent local and non-local trials from light to dark gray. b, top right. The average edge deviations (in absolute value) progressively increase when non-local interactions take place. c. While the decays of the SSN edges (purple) are consistently slower than the FID matrix (top, negative deviation), the LSN edges (yellow) vary more, and both slower and faster decays are observed (widespread deviations along the x-axis). Overall, the pattern of deviations across trials (bottom, lower triangular matrix) has a similar topography to the average FID matrix (bottom, upper triangular matrix) with Spearman’s r_s_ = 0.79. d. Standardized nodal decay times (AMI minima) of empiric trials sorted by their similarity to the trial-average (top). The trials with highest similarity show the known back-to-front gradient of timescales (bottom, left). The furthest trials from the mean display an inverse gradient (bottom, right). e. Correlation matrix of the regional AMI minima (for the nodal timescales) between all pairs of trials. The trials are sorted as in panel d. The correlation matrix displays two major blocks corresponding to the standard and inverse gradient trials. The standard-gradient trials correspond to the trials defined as local based on the edge decay-times (the strongest 10% of these are represented as blue dots in the right panel). The inverse-gradient trials (orange dots, representing approximately 10% of the total) are predominantly non-local.

We then explore the relationship between non-local interactions and the known nodal gradient of timescales (Kiebel et al., 2008; Murray et al., 2014b; Gao et al., 2020). Computing the AMI decay times of the nodal time series and sorting trials in growing order of similarity with the trial-average pattern, we recover the well-established back-to-front gradient of time-scales (Fig.3.d, bottom left). Remarkably, the trials that are the least similar to the average reveal an inverse front-to-back gradient (Fig.3.d, bottom right). The matrix of cross-trials correlations (Fig.3.e, left) shows that standard- and inverse-gradient trials are similar to themselves and anticorrelated to each other. Finally, most of the trials with an inverse front-to-back gradient are dominated by non-local higher-order interactions between regions (Fig. 3e, right; Spearman’s r_s_ = 0.42 between the degree of non-locality as observed from edge analysis and the distance from the average nodal gradient).

## Discussion

Our results reveal that the dynamics of nonlocal interactions play a key role in shaping whole brain activity. In addition to the classical back-to-front gradient of timescales, which mostly reflect local information processing, nonlocal interactions are associated with an inverse front-to-back gradient. Our work highlights the necessity to perform a trial-specific analysis since trial-averaging masks the diversity in dynamics. We propose that interactions between brain regions encode information beyond the purely nodal activity, and that such information is retained according to edge-specific characteristic lifetimes. The presence of non-local interactions in a subset of trials can be interpreted within the integration/segregation hypothesis (Shine et al.,2016), which suggests that the brain alternates moments in which the processing of information is local to moments of collective processing. The distribution of the average information decays reveals two subnetworks with short (SSN) and long (LSN) storage capability. This result is in line with multiple models (Engel et al., 2001) and experimental evidence (Buschman and Miller,2007) showing that processing of external stimuli involves (bottom-up) perception and abstraction, as well as (top-down) interpretation according to expectations (priors), embodied in the internal brain state (Friston, 2010). The presence of an inverse gradient speaks to a dialectic interaction between top-down and bottom-up processes, which remains open to interpretation while stressing once again the relevance of distributed dynamic brain processes underpinning cognition.

## Materials and Methods

### Participants

Fifty-eight right-handed and native Italian speakers were considered for the analysis. To be included in this study, all participants had to satisfy the following criteria: a) to have no significant medical illnesses and not to abuse substances or use medication that could interfere with MEG/EEG signals; b) to show no other major systemic, psychiatric, or neurological illnesses; and c) to have no evidence of focal or diffuse brain damage at routine MRI. The study protocol was approved by the local Ethics Committee. All participants gave written informed consent.

### MRI acquisition

3D T1-weighted brain volumes were acquired at 1.5 Tesla (Signa, GE Healthcare) using a 3D Magnetization-Prepared Gradient-Echo BRAVO sequence (TR/TE/TI 8.2/3.1/450 ms, voxel 1 × 1 × 1 mm3, 50% partition overlap, 324 sagittal slices covering the whole brain).

### MEG acquisition

Subjects underwent magnetoencephalographic examination in a 163 – magnetometers MEG system placed in a magnetically shielded room (AtB Biomag UG - Ulm - Germany). The preprocessing was done similarly as in (Sorrentino et al., 2018). In short, the position of four coils and of four reference points (nasion, right and left pre-auricular point and apex) were digitized before acquisition using Fastrak (Polhemus^®^). The brain activity was recorded for 7 minutes, with eyes closed, with a break at 3.5 minutes, so as to minimize the chances of drowsiness. The head position was recorded at the start of each segment. The data were sampled at 1024 Hz and a 4th order Butterworth band-pass filter was applied to select components between 0.5 and 48 Hz. During the acquisitions, electrocardiogram (ECG) and electrooculogram (EOG) were also recorded (Gross et al., 2013). These steps were done using Matlab 2019a and the Fieldtrip toolbox 2014 (Oostenveld et al., 2011).

### Preprocessing

Principal component analysis (PCA) was performed to reduce the environmental noise (Sadasivan P.K. and Narayana Dutt D., 1996; de Cheveigné and Simon, 2007). Noisy channels were removed manually through visual inspection of the whole dataset by an experienced rater. Supervised independent component analysis (ICA) was performed to eliminate the ECG (and the EOG) component from the MEG signals (Barbati et al., 2004). Trials that did not contain artefacts (either system related or physiological) or excessive environmental noise were selected.

### Source reconstruction

The data were co–registered to the native MRI. A modified spherical conductor model was used as a forward model (Nolte, 2003). The voxels in the MRI were labelled according to the Automated Anatomical Labelling (AAL) atlas (Tzourio-Mazoyer et al., 2002; Gong et al., 2009). We used the cortical regions for a total of 78 areas of interest. Subsequently, a linearly constrained minimum variance beamformer was used to compute 78 time series (one per area of interest) at a sampling frequency of 1024 Hz (Van Veen et al., 1997). Reconstructed sources were again visually inspected by an experienced rater. Of the initial 58 subjects, 44 had enough artefact-free acquisitions and were selected for further analysis. The source reconstructed signals were downsampled to 256 Hz.

### Edge-centric approach to MEG

In this work, we adopted an edge-centric approach that, rather than focusing on the local activity of the regions (nodes), represents the dynamics of the interactions between couples of brain regions (Esfahlani et al., 2020). This allowed us to characterize the whole brain network activity in terms of dynamical non-local interactions, highlighting the relational properties of each couple of nodes. Given any couple of nodes *i* and *j* and their respective source-reconstructed signals X_i_(t)and X_j_(t), we defined a characteristic time series E_ij_for the edge ij as the product of the z-scored signals, i.e.,

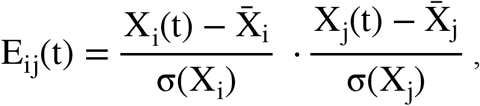

where 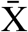 and σ(X)denote the mean and variance of the signals, respectively. One can interpret the edge co-activation time series as the unfold in time of the pairwise correlations. In fact, the average of the above expression over time corresponds to the Pearson correlation between the signals at nodes *i* and *j*. The edge time series were further analyzed by Information theoretic measures, aiming at characterizing the Information storage capability of each functional edge.

Estimation of information decay time through Mutual Information Shannon Entropy, defined as

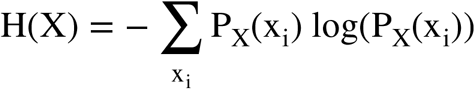

quantifies the uncertainty over the possible outcomes X_i_of a random variable X with probability distribution P_X_. If the uncertainty over future outcomes of X decreases as we measure the outcome y_i_ of another random variable Y, we conclude that X and Y represent two processes that are not independent. The new resulting uncertainty over X is then defined by

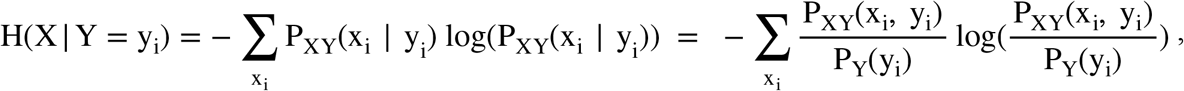

withP_XY_(X_i_, y_i_) denoting the joint probability distribution of the pair (X, Y). The weighted sum over all possible outcomes of Y defines the Conditional Entropy, i.e. the uncertainty of X conditioned on Y

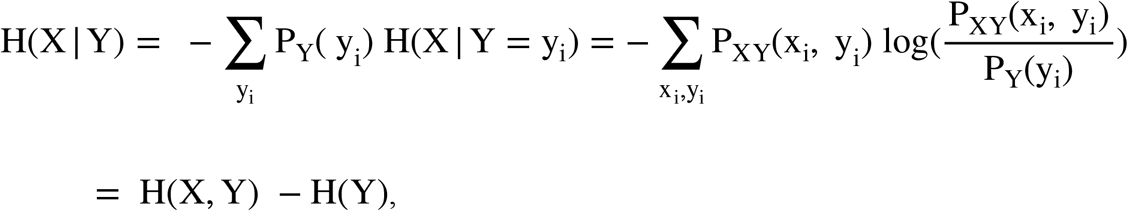

where

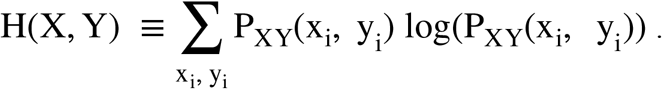

The reduction in uncertainty (or - equivalently - the increase of information) over X given by the knowledge of Y is measured by the Mutual Information (MI)

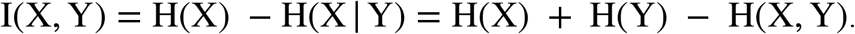

Unlike other measures, such as partial autocorrelation (PAC), Mutual Information (MI) statistical dependencies which take into account nonlinear interactions, which are ubiquitously found in brain data (Paluš, 1996; Stam, 2005)P. In order to quantify the time span before the information in a signal X(t)is lost, we rely on the Auto Mutual Information (AMI) i.e., the MI between the signal and its time delayed copy Y = X(t – τ). According to previous works on M/EEG (Jeong et al., 2001; Gómez et al., 2007), a stable estimate of the probability distribution of any real-valued signal X(t) is obtained by dividing the data into 64 bins. The joint probability distribution for the pair (X(t), X(t – τ)), needed for the evaluation of the AMI, is given by a 64×64 contingency matrix which is the joint histogram of the two variables. The AMI decays as a function of the delay τ, from a maximal value at τ= 0 to a relatively stable lower value as τ→∞. The more gentle (“slower”) the decay, the longer the information lasts in the signal. It should be mentioned that there exists no unique estimator for information storage. We chose AMI since we were interested in an interval estimate, rather than a point estimate (see for example (Wibral et al., 2014)). The same algorithm was used to compute the nodal decay-times based on the nodal time series.

### Information Storage Capability of the functional edges

For each co-activation signal E_ij_, we estimated the AMI profile (Fig.1.b) and we evaluated the time delay τcorresponding to the AMI reaching a stable minimum i.e., when the original signal and its τ-delayed versions have become maximally independent. To do so, we first fit the last 80 points of every AMI profile to a straight line, so as to find the stable minimum. Then, we found the τ corresponding to the moment where the AMI decay falls within a threshold, defined as 1 standard deviation from the stable minimum. Examples of the estimate of the AMI minimum for different edges, at the single epoch level, are shown in Supplementary figure S2. An analysis for different thresholds (number of standard deviations around the stable minimum) is found in the supplementary material (Fig.S3). Averaging across 10 time windows (epochs) of 10 seconds and across 44 subjects, we found the N × N Functional Information Decay (FID) matrix (where N = 78 is the number of brain regions), containing the decay times for each edge (Fig.1.d). The AMI analysis of the co-activations shows that the decay times are different among edges, as revealed by the histogram in Figure 2.b (blue). Selecting the edges from either the left or right tails of the distribution leads to the appearance of two topographically organized subnetworks (Fig.2.a).

### Surrogate analysis

#### Leakage analysis

We designed surrogate analysis to exclude that linear mixing alone might spuriously generate the patterns observed in the FID matrix. To this end, we generated for each subject n Gaussian time-series, with n = number of regions. Then, the subject-specific leadfield matrix was used to reconstruct the sensor signals for each subject. White noise correlated as 1/sensor distance, was added to the sensor time series with SNR = 12. Following this step, the sensor time series were inverted and new surrogate source-level time series were generated. On these source-level surrogates, we have computed the edge time-series and the FID matrix as described previously.

#### Time-shuffled and phase-randomized surrogates

Firstly, we sought to investigate if the observed decay times might be derived by the spectral nodal properties alone. To this end, we generated n gaussian processes, with n = number of regions, we fourier-transformed them, and we multiplied the resulting power spectra by the power spectra of the observed time-series. Finally, we anti transformed and obtained surrogate time-series that are independent gaussian processes endowed with the same spectral power as the original data (G-surrogates). Secondly, we sought to investigate if the time decays convey a dynamical feature of the time-series or, alternatively, if they can be explained by static correlations. Hence, starting from the original data, we generated surrogates preserving not only the nodal spectral properties, but also the cross-spectrum (static functional connectivity). To this end, we shifted by a random phase (extracted from a unimodal distribution) each frequency of the Fourier transformed nodal signals. The same shift was uniformly applied to each region. Hence, we obtained new surrogates that preserve the functional connectivity while not showing the dynamic of the original data.

## Acknowledgments

The authors thank Michele Allegra for insightful discussions. The work was supported by the University of Naples Parthenope “Ricerca locale” grant, by the grant ANR-17-CE37-0001 - CONNECTOME, by the European Union’s Horizon 2020 research and innovation programme under grant agreement No. 945539 (SGA3) Human Brain Project and VirtualBrainCloud No. 826421.

## Supplementary Material

**Fig. S1.**
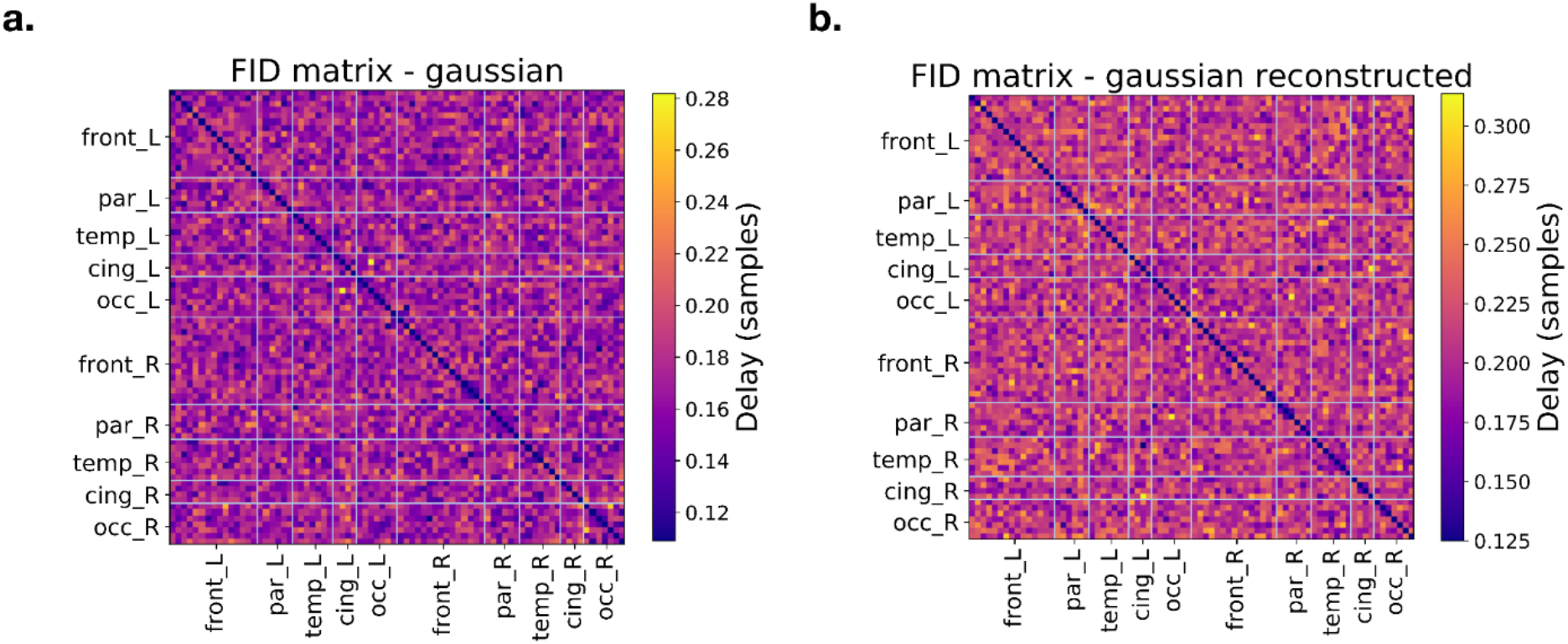
Leakage analysis pipeline. a. FID matrix computed from Gaussian surrogates. The decay times were estimated from random surrogates analogously to the procedure used with the original data. Hence, rows and columns represent brain regions, while matrix entries are the corresponding decay minima. b. Starting from the Gaussian surrogates to the left, the surrogates were outprojected, for each subject, using the corresponding mixing matrix, as to obtain new sensor-level surrogates. White noise, correlated as 1/distance between sensors, was added to the sensor-level surrogate with a SNR =12. The sensor-level surrogates were then inverted and new source-level data were generated. The minima of the decay of the AMI were the recomputed on the newly obtained source-level surrogates, to check that the spatial mixing of otherwise independent processes did not introduce the spatial patterns observed in the empirical data.

**Fig. S2:**
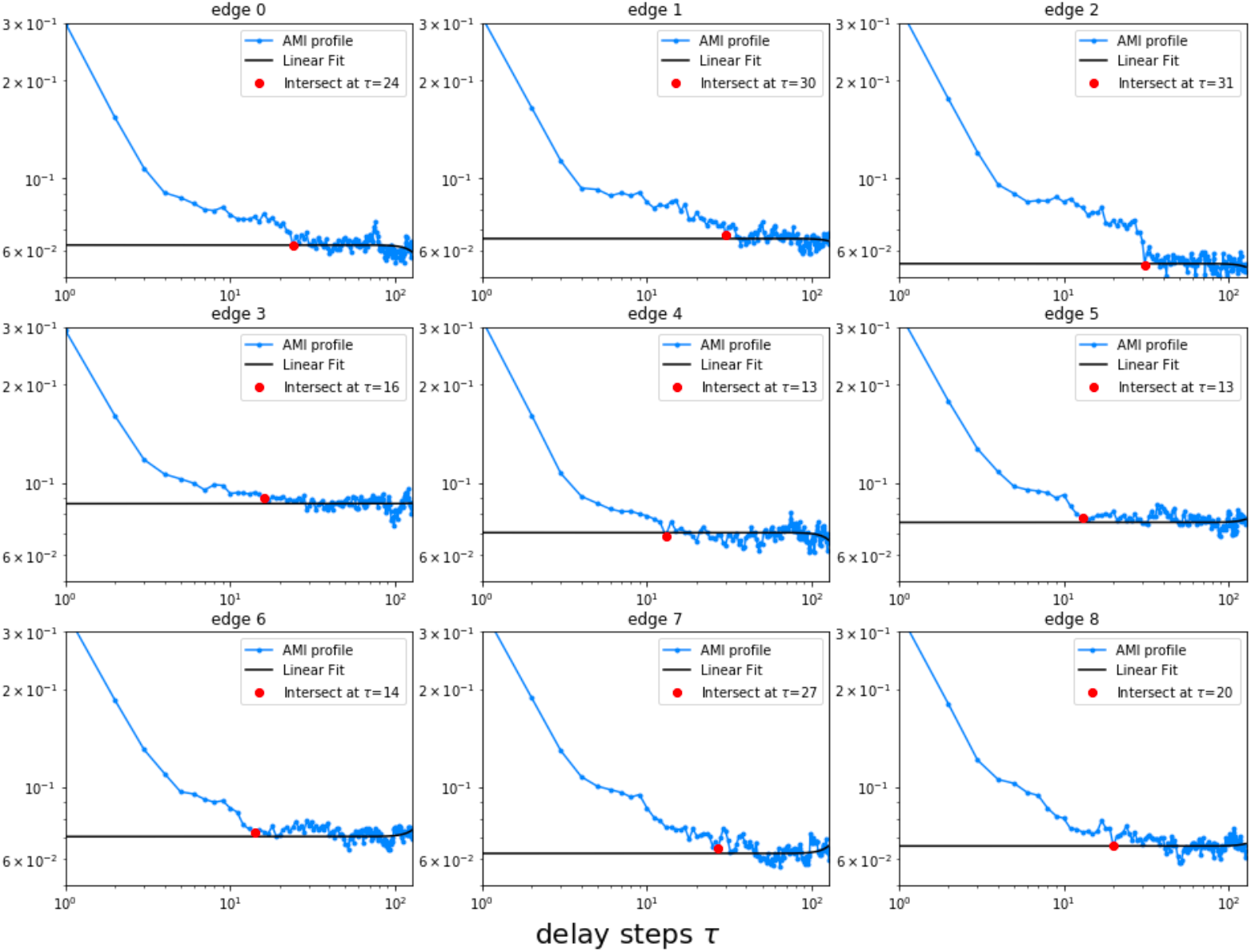
Examples of the AMI decays of 9 randomly selected edges in a single trial (i.e., form the same 10 seconds-long epoch from a randomly selected subject) (log-log scale). In order to estimate the time after which the co-activation time series and its delayed version become maximally independent, we fit the last 80 points of the AMI profile (blue) to a straight line (black) and we find the first point (red) where the difference between the AMI decay and the fitted line is below a threshold. The results reported in the main text refer to the threshold set to 1 standard deviation from the AMI tail.

**Fig.S3:**
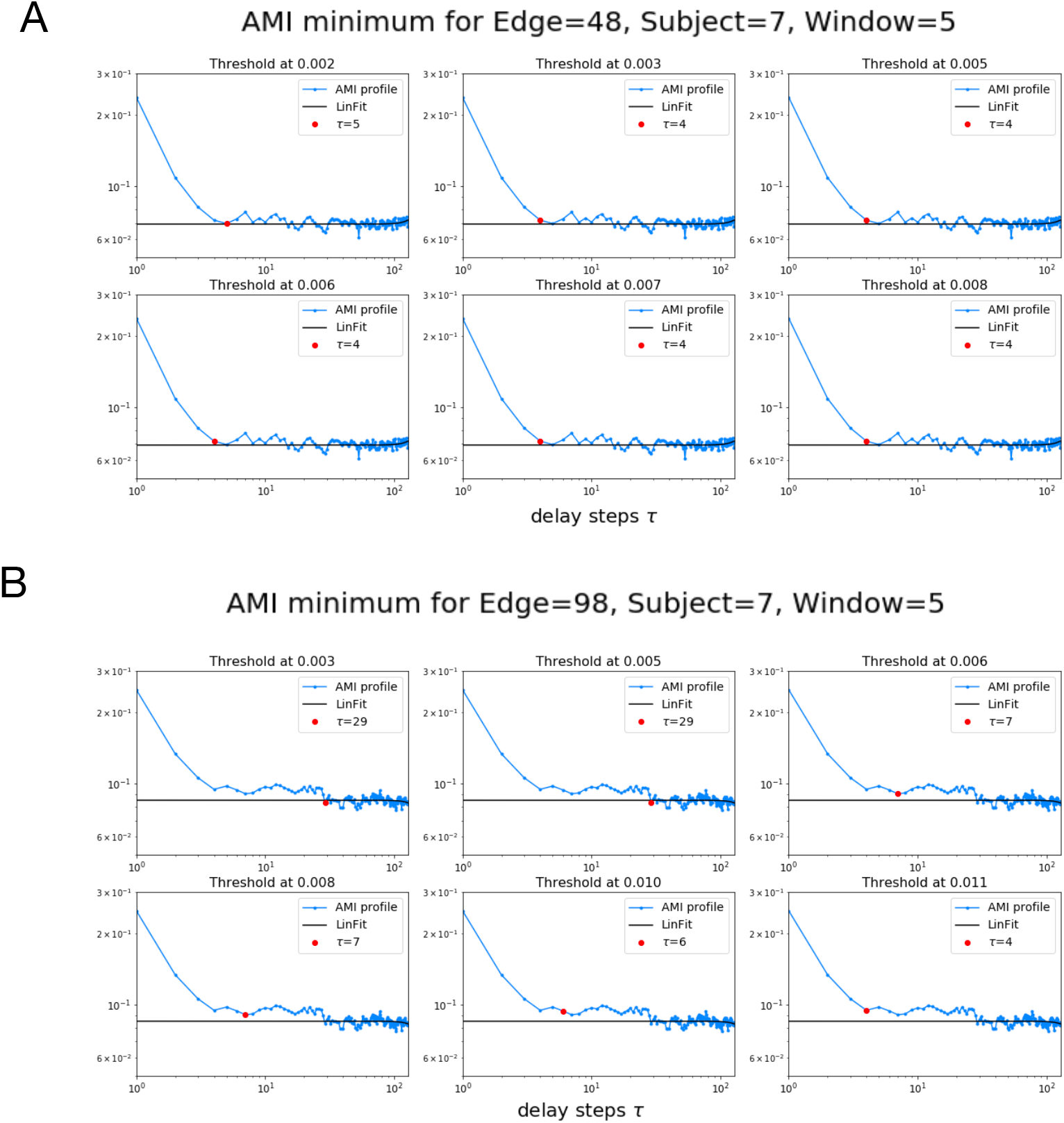
The blue lines show the AMI decay profile for a single epoch. The red dots represent the estimate of the decay-times. The decay-time is measured as the first time that the AMI profile approaches the null-line (fitted to the tail of the AMI distribution, in black). The distance from the null-line, considered as a threshold, is defined as multiples of the standard deviation ([1, 1.5, 2., 2.5, 3., 3.5]) of the AMI tail. The six panels show the corresponding estimate of the decay time as the threshold is increased, from top-left to bottom right. (A) An example is drawn from a group of edges where the AMI decay profile approaches the minimum directly. The corresponding estimate of the decay time is stable across thresholds. (B) An example is drawn from a subset of edges where the decay profile does not approach the minimum directly. In such cases, the AMI decay profile shows a fat tail before reaching a stable minimum. Since we are interested in the time required for the information to be completely extinguished in an edge, we considered the smallest threshold for the estimation of all the minima. This provides a stable estimate of the minimum over all the observed decay profiles.

### Detailed analysis of the average edge topography

The SSN and LSN subnetworks are topographically distinct, as shown in Fig.2.a. Edges with short information storage are often incident upon occipital regions, although not uniformly. In fact, it appears that the cunei and the occipital gyri are specifically involved in the SSN. These regions roughly correspond to Brodmann areas 17, 18 and 19, and are known to be hierarchically related to the extraction of abstract shapes from the visual input. The cuneus directly receives inputs from the retina, that are subsequently processed up to the superior occipital gyrus, in the visual associative cortex, which is responsible of the perception of the properties of objects and plays a role, for instance, in face and object recognition (Renier et al., 2010; Freud et al., 2017). The occipital regions are connected bilaterally to the inferior parietal gyrus (e.g. the angular and supramarginal gyri as well as the inferior parietal lobe), the posterior temporal lobes and frontal/ prefrontal cortex. The posterior part of the parietal lobe has been included in the “dorsal stream” of the visual system, which is mainly concerned with the analysis of spatial relationships of objects, and the information about the position of the body in space (Goodale and Milner, 1992). With regard to the temporal lobe, the posterior part appears to be selectively involved in the SSN. From a structural standpoint, the inferior-longitudinal fasciculus (ILF), as well as an “indirect” stream of U-bundle fibers, connects the occipital and temporal areas (Herbet et al.,2018). Functionally, these regions are related to either the high-level analysis of images, such as the inferior temporal lobe, whose lesion compromises the ability to recognize facial expressions, or to the perception of- and attendance to auditory stimuli, as in the case of the superior temporal lobe (Vander Ghinst et al., 2016). Finally, in the frontal lobes, the SSN appears to involve the ventrolateral and dorsolateral prefrontal cortices, which participate to higher functions such as allocation of attention, and are typically considered the end of the ventral and dorsal streams, respectively (Goldman and Rosvold, 1970; de Pasquale et al., 2012). In short, all the regions that are known to be relevant in the process of sensory stimuli emerge as a temporally homogeneous fast network carrying information from visual and acoustic areas, through layers of abstraction, up to areas related to conscious perception.

The opposite network, to which we refer to as the long storage network (LSN), clusters topographically in frontal and temporal regions. The regions involved in the LSN appear to be linked anatomically by the uncinate fasciculus. In fact, the LSN mainly connects the orbitofrontal cortex within the frontal cortex to the temporal poles in the temporal lobes. Patients who underwent brain surgery involving the removal of the uncinate fasciculus showed impairment of verbal memory, naming of objects, verbal fluency, and name anomia. These symptoms did not appear in patients whose surgery did not encompass the removal of the connections between the orbitofrontal cortex and the temporal lobes (Papagno et al., 2011). This evidence might highlight the functional significance of the LSN for the integration of information across these anatomically linked regions. Furthermore, the semantic variant of primary progressive aphasia and herpes encephalitis, both diseases that induce a damage to the temporal poles, cause disruptions in categorical discrimination, word comprehension and naming (Noppeney et al., 2007). Recently, Warren et al. (Warren et al., 2009), using fMRI showed that, during narrative speech comprehension, the left anterior basal temporal cortex shows high correlation with the left anterior basal frontal cortex, the left anterior inferior temporal gyrus as well as with the corresponding homotopic temporal cortex contralaterally. From a neurophysiological perspective, the N400 response is an event-related potential (ERP) evoked ~400 ms after (potentially) meaningful material is presented. Using MEG, the N400 was localized in the superior temporal sulcus (Marinkovic et al., 2003), and intracranial recordings showed it originates in the anteroventral temporal lobe. Furthermore, the SSN entails more long-range inter-hemispheric connections as compared to the LSN, which might suggest that interhemispheric coordination is specifically achieved via fast interactions (perhaps specifically relying on fast, white-matter bundles such as the corpus callosum).

